# An atypical CRISPR-Cas locus in *Symbiobacterium thermophilum* flanked by a transposase, a reverse transcriptase, the endonuclease MutS2 and a putative Cas9-like protein

**DOI:** 10.1101/252296

**Authors:** Sandeep Chakraborty

## Abstract

Clustered regularly interspaced short palindromic repeats (CRISPR) is a prokaryotic adaptive defense system that assimilates short sequences of invading genomes (spacers) within repeats, and uses nearby effector proteins (Cas), one of which is an endonuclease (Cas9), to cleave homologous nucleic acid during future infections from the same or closely related organisms. Here, a novel CRISPR locus with uncharacterized Cas proteins, is reported in *Symbiobacterium thermophilum* (Accid:NC 006177.1) around loc.1248561. Credence to this assertion is provided by four arguments. First, the presence of an exact repeat (CACGTGGGGTTCGGGTCGGACTG, 23 nucleotides) occurs eight times encompassing fragments about 83 nucleotides long. Second, comparison to a known CRISPR-Cas locus in the same organism (loc.355482) with an endonuclease Cas3 (WP 011194444.1, 729 aa) ∼10000 nt upstream shows the presence of a known MutS2 endonuclease (WP 011195247.1, 801 aa) in approximately the same distance in loc.1248561. Thirdly, and remarkably, an uncharacterized protein (1357 aa) long is uncannily close in length to known Cas9 proteins (1368 for *Streptococcus pyogenes*). Lastly, the presence of transposases and reverse transcriptase (RT) downstream of the repeat indicates this is one of an enigmatic RT-CRISPR locus, Also, the MutS2 endonuclease is not characterized as a CRISPR-endonuclease to the best of my knowledge. Interestingly, this locus was not among the four loci (three confirmed, one probable) reported by crisperfinder (http://crispr.i2bc.paris-saclay.fr/Server), indicating that the search algorithm needs to be revisited. This finding begs the question ‐ how many such CRISPR-Cas loci and Cas9-like proteins lie undiscovered within bacterial genomes?

## Introduction

The seemingly innocuous discovery of clustered regularly interspaced short palindromic repeats (CRISPR) in bacteria [1] was later established as a prokaryotic adaptive defense system that memorizes short sequences of invading genomes (spacers) within repeats [2–4], and uses nearby effector proteins (Cas), one of which is an endonuclease (Cas9) [5–7], to cleave subsequent infections from the same organism [8,9].

The CRISPR-Cas system is divided into two classes ‐ class 1 (90%) and class 2 (10%) ‐ based on whether there are multiple effectors or a single effector, respectively [10,11]. The simplicity of the single effector class 2 CRISPR-Cas systems has fueled an incredible acceleration in gene-editing technologies that have raised prospects of human disease management [12–14], simultaneously raising serious ethical issues [15–17].

Technical limitations of the application of the CRISPR-Cas system for therapeutic purposes arise from the non-specificity of cleavage (off-targets) [18,19], with several methods available to verify such non-desirable cleavages [20,21]. Understanding the structural basis of target recognition in these Cas9 proteins [22] helps in generating Cas9 variants through amino-acid substitutions that moderate target site recognition and cleavage to minimize off-target cleavage [23]. It has recently been proposed that there are ‘pre-existing humoral and cell-mediated adaptive immune responses to Cas9 in humans’ [24]. In addition to generating Cas9 variants that do no elicit immune response, unearthing novel Cas9 proteins could be another strategy to circumvent this problem.

Bacterial sequence diversity is mind-boggling. Even in reasonably well-characterized species, like *Symbiobacterium thermophilum* [25,26], there are many proteins that are uncharacterized. Could it be that some of the proteins are CRISPR-Cas related? Here, the genome of *S. thermophilum* has been analyzed for such proteins.

## Results and discussion

There are four reasons that strongly support the assertion that loc.1248561 is a novel CRISPR-Cas locus.

1. An exact repeat ‐ CACGTGGGGTTCGGGTCGGACTG, 23 nucleotides ‐ occurs eight times, and has integrated seven fragments that are ∼83 nucleotides long (Fig 1a), and is predicted (RNAfold [27]) to have the proper folding to ensure cleavage.

**Figure 1:**
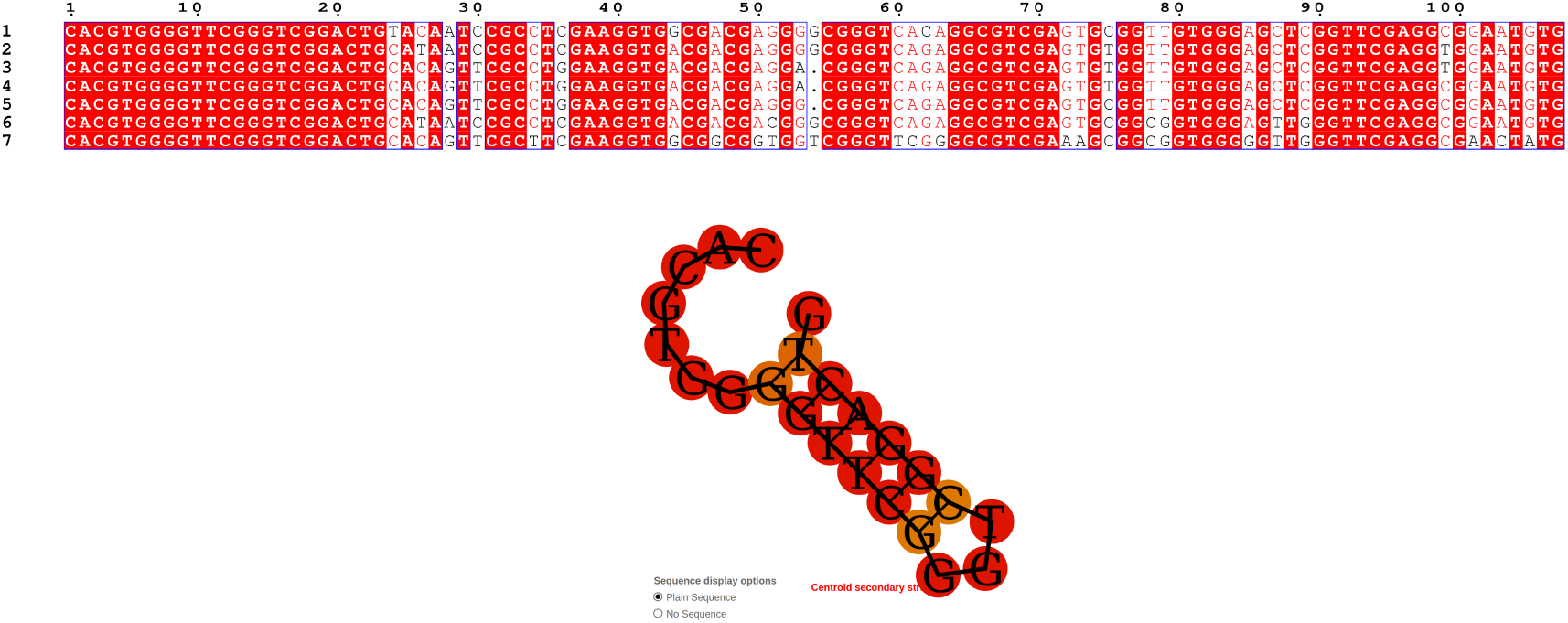
Repeats and spacers in the new locus: The sequence CACGTGGGGTTCGGGTCGGACTG (23 nt) is repeated eight times, and encompasses seven fragments about 83 nucleotides long. The secondary structure of the crna is predicted by RNAfold [27]. Note there two mismatched pairs (G-T). The cleavage site is predicted to the sequence at the end of the stem on the 3’ end [36].
2. A known CRISPR-Cas locus in the same organism (loc.355482) has a 729 aa endonuclease CAS3 (WP 011194444.1) at 9440 nt from the repeat location (Table 1). Analogously, loc.1248561 has a 801 aa endonuclease (MutS2) 10819 nt away from the repeat location.
3. An uncharacterized protein (1357 aa) long located between the MutS2 endonuclease and the repeats is uncannily close in length to known Cas9 proteins (1368 aa for *Streptococcus pyogenes*).

**Table 1:** Annotation of open-reading frames (ORF) around the known CRISPR-loci (loc.355482, KNOWN) and the new CRISPR-loci (loc.1248561, NEW) in *Symbiobacterium thermophilum*: ORF numbering is based on the ORFs obtained using ‘getorf’ (SI ORFS.fa). ORFs that are 15000 nt from the loci are annotated on both sides (pre and post). The distance (9440) of the helicase/endonuclease CAS3 (WP 011194444.1, 729 aa) in loc.355482 to the repeats is almost the same as the distance (10819) of MutS2 endonuclease (WP 011195247.1, 801) to the repeats in loc.1248561. None of the apparently bona-fide proteins between the MutS2 and the repeats, some having ORFs as long as 1357 aa, are characterized. Intriguingly, the 1357 is uncannily close to length of known Cas9 proteins (1368 for *Streptococcus pyogenes*) ‐ *Symbiobacterium thermophilum* does not have a Cas9 CRISPR-Cas locus. Finally, a group II intron reverse transcriptase (RT) downstream of the repeat suggest that this locus is from a family of enigmatic RT-CRISPR systems [29], while the locus also has transposases that explain the CRISPR-Cas genesis.
4. The presence of a reverse transcriptase (RT, WP 011194877.1) downstream of the repeat indicates this locus is a RT-CRISPR that integrates RNA (after the RT transcribes it into DNA) [28–30], possibly using a proximal transposase [31,32].

| Dir | ORF | LEN | Annotation | Start | End |
| --- | --- | --- | --- | --- | --- |
| KNOWN-pre | 2548 | 703 | ? | 340008 | 342116 |
|  | 2559 | 444 | IS110 family transposase | 342291 | 343622 |
|  | 2590 | 442 | IS256 family transposase | 345631 | 346956 |
|  | 2611 | 729 | CAS helicase/endonuclease CAS3 | 347066 | 349252 |
|  | 2633 | 625 | TYPE I-C CAS8C/CSD1 | 349868 | 351742 |
|  | 2640 | 385 | TYPE I-C CAS7/CSD2 | 351697 | 352851 |
|  | 2656 | 392 | IS5/IS1182 family transposase | 354046 | 355221 |
|  |  |  | Repeat location 355482 |  |  |
| KNOWN-post | 2689 | 683 | ? | 357654 | 359702 |
|  | 2739 | 695 | Hypothetical protein | 363051 | 365135 |
|  | 2748 | 308 | Hypothetical protein | 365125 | 366048 |
|  | 2774 | 302 | Hypothetical protein | 368033 | 368938 |
| NEW-pre | 8862 | 646 | ABC Transporter ATP-binding protein | 1234429 | 1236384 |
|  | 8867 | 211 | DUF421 domain-containing protein | 1235825 | 1237183 |
|  | 8871 | 320 | Hypothetical protein | 1236972 | 1238081 |
|  | 8886 | 801 | Endonuclease MutS2 | 1238171 | 1240573 |
|  | 8902 | 215 | ? | 1242464 | 1243420 |
|  | 8913 | 1357 | ? (Putative Cas9) | 1240660 | 1244943 |
|  |  |  | Repeat location 1248561 |  |  |
| NEW-post | 8950 | 305 | ? | 1249228 | 1250463 |
|  | 8974 | 264 | ? | 1252673 | 1253659 |
|  | 8981 | 409 | Tyrosine-trna ligase | 1253070 | 1254419 |
|  | 8988 | 325 | ? | 1254535 | 1255509 |
|  | 9007 | 611 | Group II intron Reverse transcriptase/maturase | 1255638 | 1257593 |
|  | 9015 | 254 | Adenylyltransferase | 1257668 | 1258747 |
|  | 9020 | 269 | Hypothetical protein | 1258183 | 1259478 |
|  | 9033 | 530 | Transposase | 1259556 | 1261148 |
|  | 9054 | 408 | Transposase | 1262612 | 1263895 |

### The 23 nt repeats

The CRISPR repeats are typically varying in size from 23 to 47nt [33], which are transcribed and cleaved into mature CRISPR RNAs (crRNA) [34]. Although, some CRISPR-Cas loci have an trans-activating CRISPR RNA molecule (tracrRNA) that co-purifies with Cas9, and is essential for DNA interference [35], the locus here seems to have has no such sequence (the tracrRNA is homologous, but not exact to the crna). The secondary structure of the crna (Fig 1b) predicted by RNAfold [27] shows analogous folding to known repeats that allows cleavage [36].

Interestingly, this locus was not among the four loci (three confirmed, one probable) reported by Crisperfinder [33]. PILER-CR also did not detect this locus [37]. CRISPRdigger [38] and CRT [39] have not been evaluated. Crisperfinder has several criteria for choosing repeats and spacers, and ‘and the percentage of identity between spacers is not allowed to exceed 60%’. The presence of related protein in the vicinity of repeats (as well as it folding properties) are much stronger criteria, that would override such homology thresholds. This indicates that those parameters need to be revisited.

### MutS2 endonuclease

To the best of my knowledge, MutS2 is not characterized as a CRISPR-endonuclease. MutS2 from *Thermus thermophilus* was found to suppress homologous recombination [40], whereas a recent study in *Bacillus subtilis* demonstrated the opposite phenomenon [41]. Also, though MutS2 was mentioned in the protocol for ‘CRISPR/Cas9 Editing of the Bacillus subtilis Genome’, there is no indication that its CRISPR-like endonuclease activity was discussed. Another study emphasized the requirement of all domains of this protein for its functionality [42]. The considerable sequence homology to other MutS2 proteins expectedly translates into significant structural homology (Fig 2a) on a predicted structures, and will help in identifying domains/active sites [43] and other promiscuous functions, if they exist [44].

**Figure 2:**
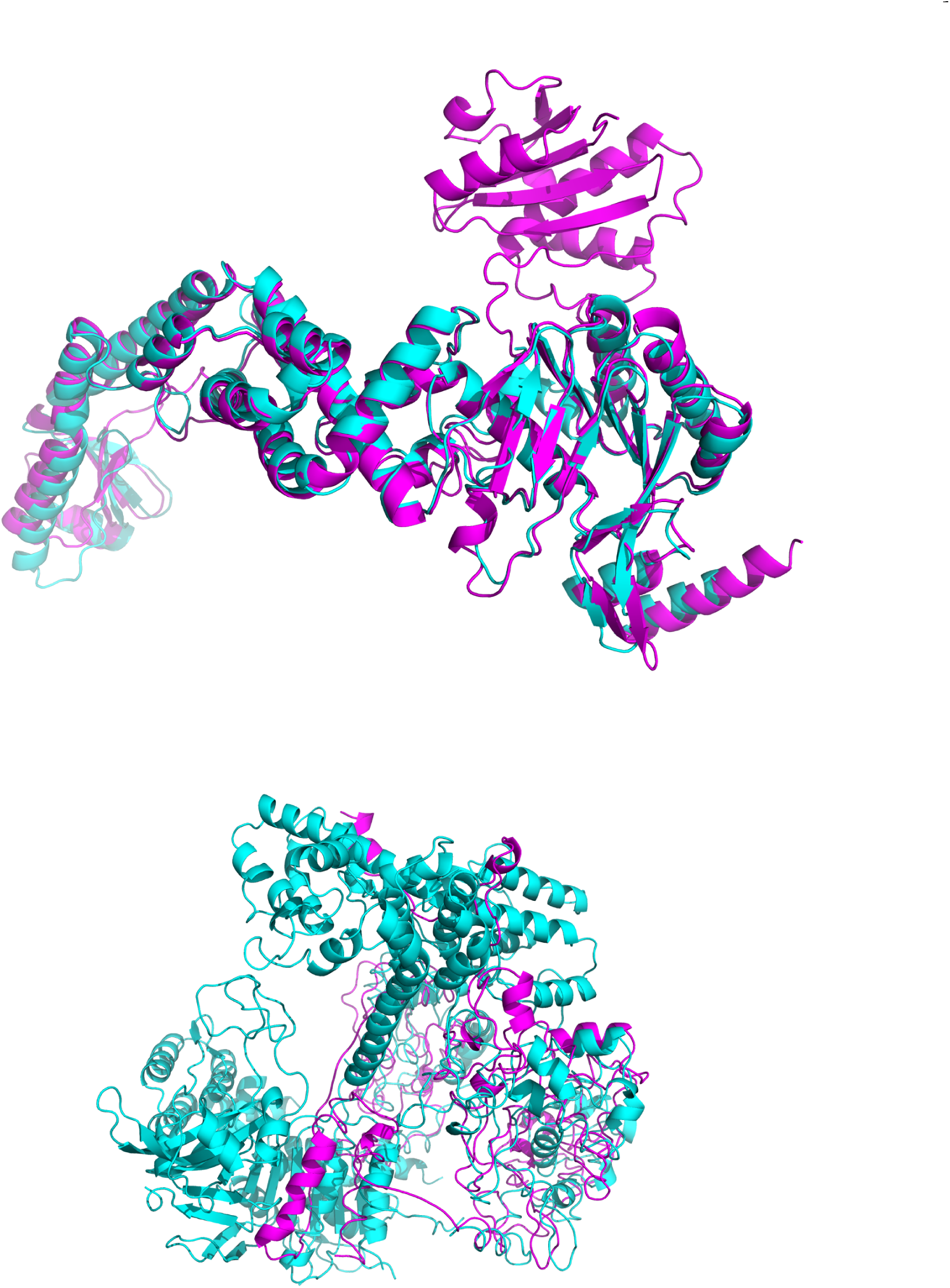
Structure prediction and superimposition for Muts2 and the novel Cas9: (a) Structure of Muts2 (cyan) predicted by SWISSMODEL using PDBid:5AKC (magenta) (‘MutS in complex with the N-terminal domain of MutL’ from *Escherichia coli* [55]) as template. The superimposition shows good structural homology. (b) Structure of novel Cas9 (magenta) predicted by SWISSMODEL using PDBid:4OO8 (cyan) (‘Crystal structure of *Streptococcus pyogenes* Cas9 in complex with guide RNA and target DNA’ [56]) as template. Due to lack of sequence homology, the structure prediction method does not predict a reasonable structure. Future functional and structural characterization of this putative Cas9 will reveal these features.

### Reverse transcriptase based CRISPR loci

The reverse flow of genetic information from RNA to DNA, essential to the retroviral replication cycle [45], was a momentous discovery (made independently by Temin and Baltimore in 1970) challenging Crick’s famous ‘central dogma’ [46] that information flowed from DNA to RNA to proteins, and never in the reverse direction. These enzymes were identified in bacteria almost two decades later [47]. In many Type III CRISPR systems, a reverse transcriptase (RT) is fused to a Cas1, which in the marine bacterium *Marinomonas mediterranea*, was shown to mediate the *in vivo* acquisition of RNA spacers [28]. Two recent studies looked at the evolutionary origins of RT-CRISPR, and looked mostly at RT-Cas1 fusions [29, 30]. Moreover, these work seeded on known CRISPR-Cas loci, and thus would not have looked at the locus described here. The spacer sequences here (Fig 1a) could not identify any known organism, echoing the observation that ‘the vast majority of RT-linked CRISPR spacers come from a distinct sequence pool, the nature of which remains enigmatic’ [29].

### Putative Cas9-like protein

Another fascinating observation is the presence of an uncharacterized open reading frame (ORF, 1357 aa long), which is uncannily close to the size of known Cas9 proteins (1368 for *Streptococcus pyogenes*). Due to the lack of homology with known Cas9, the structure prediction (using SWISS-MODEL [48]) did not result in any realistic structure (Fig 2b), in contrast to the the MutS2. There is another long ORF (>200 aa) between the MutS2 endonuclease and the repeat location, and three such ORFs after the repeat location. These are all uncharacterized, even on a complete BLAST [49] to the nr database.

### Conclusion and future work

Seminal work on the enumeration and classification of CRISP-Cas systems first used the highly conserved Cas1 as the seed [10], and followed it up by using known CRISP-Cas sequences from databases [11] in an iterative search method. However, this underestimates bacterial genome diversity ‐ and a completely novel CRISPR locus will escape detection. On, the other hand using repeated sequences as the starting point, and trying to look for relevant proteins (endonucleases, transposases, reverse transcriptases) could identify more CRISPR-Cas loci, and related Cas proteins. Even, the presence of uncharacterized protein coding open reading frames of reasonable length and number should suffice to tag a repeat as a putative CRISPR-Cas loci. Future work will build such a database.

## Materials and methods

ftp://ftp.ncbi.nih.gov/genomes/refseq/bacteria/Symbiobacterium thermophilum/latest assembly versions/ provided the *Symbiobacterium thermophilum* genome and annotation. The getorf program from the EMBOSS suite [13] was used to obtain the ORFs (SI ORFS.fa). ORFs that are 15000 nt from the loci are annotated on both sides using YeATS [50–52] based on the Ncbi annotation.

RNAfold [27] was used to predict the secondary structure of the RNA repeat. Structures of the MutS2 and the novel predicted Cas9 were predicted using SWISS-MODEL [48]. Structures were superimposed using MUSTANG [53]. MSA figures were generated using the ENDscript server [54]. Protein structures were rendered by PyMol (http://www.pymol.org).

